# A structural mechanism for directing inverse agonism of PPARγ

**DOI:** 10.1101/245852

**Authors:** Richard Brust, Jinsai Shang, Jakob Fuhrmann, Jared Bass, Andrew Cano, Zahra Heidari, Ian M. Chrisman, Anne-Laure Blayo, R. Patrick Griffin, Theodore M. Kamenecka, Travis S. Hughes, Douglas Kojetin

## Abstract

Small chemical modifications can have significant effects on ligand efficacy and receptor activity, but the underlying structural mechanisms can be difficult to predict from static crystal structures alone. Here we show how a simple phenyl-to-pyridyl substitution between two common covalent orthosteric ligands targeting peroxisome proliferator-activated receptor gamma (PPARγ) converts a transcriptionally neutral antagonist (GW9662) into an inverse agonist (T0070907). X-ray crystallography, molecular dynamics simulations, and mutagenesis coupled to activity assays reveal a water-mediated hydrogen bond network linking the T0070907 pyridyl group to Arg288 that is essential for inverse agonism. NMR spectroscopy reveals that PPARγ exchanges between two long-lived conformations when bound to T0070907 but not GW9662, including a conformation that prepopulates a corepressor-bound state, priming PPARγ for high affinity corepressor binding. Our findings demonstrate that ligand engagement of Arg288 may provide new routes for developing PPARγ inverse agonist.

## Introduction

The nuclear receptor peroxisome proliferator-activated receptor gamma (PPARγ) is a molecular target for insulin sensitizing drugs, including the thiazolidinedione (TZD) or glitazone class of antidiabetic drugs ^1^. TZDs are full agonists of PPARγ that induce transcriptional activation resulting in the differentiation of multipotent mesenchymal stem cells (MSCs) into adipocytes or fat cells. Unfortunately, PPARγ agonists used as therapeutic agents in patients with type 2 diabetes mellitus (T2DM) display adverse side effects, including differentiation of bone tissue into fat resulting in brittle bone.

Although originally it was thought that full activation of PPARγ was required for antidiabetic efficacy, recent studies have shown that antidiabetic PPARγ ligands can span a wide range of efficacies—including full and partial agonists, antagonists, and inverse agonists that have robust or mild activating, neutral, or repressive transcriptional properties, respectively ^2-5^. Importantly, repressive PPARγ modulators decrease fat accumulation in bone and promote bone formation ^5,6^, and pharmacological repression or antagonism of PPARγ is implicated in the treatment of obesity ^7,8^ and cancer ^9-11^.

In order to determine how to promote all of the positive effects of pharmacologically targeting PPARγ on antidiabetic efficacy, bone formation, anti-obesity, and cancer treatment, we need to understand the structural mechanisms that elicit PPARγ activation (agonism) and repression (inverse agonism). These distinct pharmacological phenotypes of PPARγ ligands are dictated by ligand-dependent recruitment of transcriptional coregulator proteins (coactivators and corepressors) to the PPARγ ligand-binding domain (LBD). The LBD contains the orthosteric ligand-binding pocket for endogenous ligands, which is also the binding site for most synthetic ligands, and the activation function-2 (AF-2) coregulator binding surface. The AF-2 surface is composed of three LBD structural elements: helix 3, helix 5, and the critical helix 12 that dynamically moves between two or more conformations in the absence of ligand ^12^.

The structural mechanisms affording activation of PPARγ are well understood. Agonists stabilize an active state of the AF-2 surface by forming hydrogen bonds with residues near helix 12. Full agonists form a critical hydrogen bond with the phenolic side chain of Y473 on helix 12, strengthen coactivator and weaken corepressor binding affinities, respectively, which induces robust transcriptional activation ^12-14^. Partial or graded agonists do not hydrogen bond to Y473, but mildly stabilize helix 12 via interactions with other regions of the ligand-binding pocket, resulting less pronounced changes in coregulator affinity and transcriptional activation ^13-15^. Neutral or passive antagonists, which make unfavorable interactions with F282 on helix 3, do not stabilize helix 12 and display negligible changes in activation ^3^. These findings have established the structural mechanisms for eliciting robust (agonist), weak (partial agonist), or no (antagonist) transcriptional activation of PPARγ. An inverse agonist has a profile opposite of an agonist, increasing the binding affinity of corepressors and decreasing the binding affinity of coactivators resulting in transcriptional repression. However, relatively few studies have explored the structural mechanisms by which ligands repress PPARγ transcription ^6,16^, and it remains poorly understood how to design inverse agonists.

Here we compare two commonly used covalent PPARγ ligands, GW9662 ^17^ and T0070907 ^18^, which are referred to as antagonists not because of their effects on PPARγ transcription, but because they covalently attach to C285 and physically block ligand binding to the orthosteric ligand-binding pocket. Remarkably, despite differing only by a simple methine (CH) to nitrogen substitution, T0070907 displays properties of an inverse agonist compared to GW9662, which shows negligible effects on transcription ^17-19^. To understand the structural basis for these effects, we solved the crystal structure of T0070907-bound PPARγ, which revealed no major overall structural differences compared to a crystal structure of GW9662-bound PPARγ that would explain the difference in efficacy. However, detailed structural analysis revealed a water-mediated hydrogen bond network that uniquely links R288 to the T0070907 pyridyl group—an interaction that cannot occur with GW9662, which lacks a hydrogen bond acceptor. NMR analysis revealed that T0070907-bound PPARγ populates two long-lived structural conformations, one of which resembles the state populated by GW9662 and a unique state that is similar to the corepressor-bound state, revealing a novel structural mechanism for directing inverse agonism of PPARγ.

## Results

### T0070907 is an inverse agonist

GW9662 and T0070907 (Figure 1A) contain the same 2-chloro-5-nitro-*N*-phenylbenzamide scaffold but differ by a simple atom change: a ring carbon in GW9662 phenyl group is replaced by a nitrogen-containing pyridyl group in T0070907. Using a cell-based transcription assay (Figure 1B), we compared GW9662 and T0070907 to other noncovalent activating and repressive PPARγ compounds (Figure S1). T0070907 repressed PPARγ transcription relative to DMSO treated cells, opposite to the agonist rosiglitazone that increased PPARγ transcription, whereas GW9662 did not significantly affect PPARγ transcription. The repressive efficacy of T0070907 is similar to or better than SR10221 and SR2595, respectively, which are analogs of the neutral antagonist/nonagonist parent compound SR1664 ^6^.

**Figure 1.**
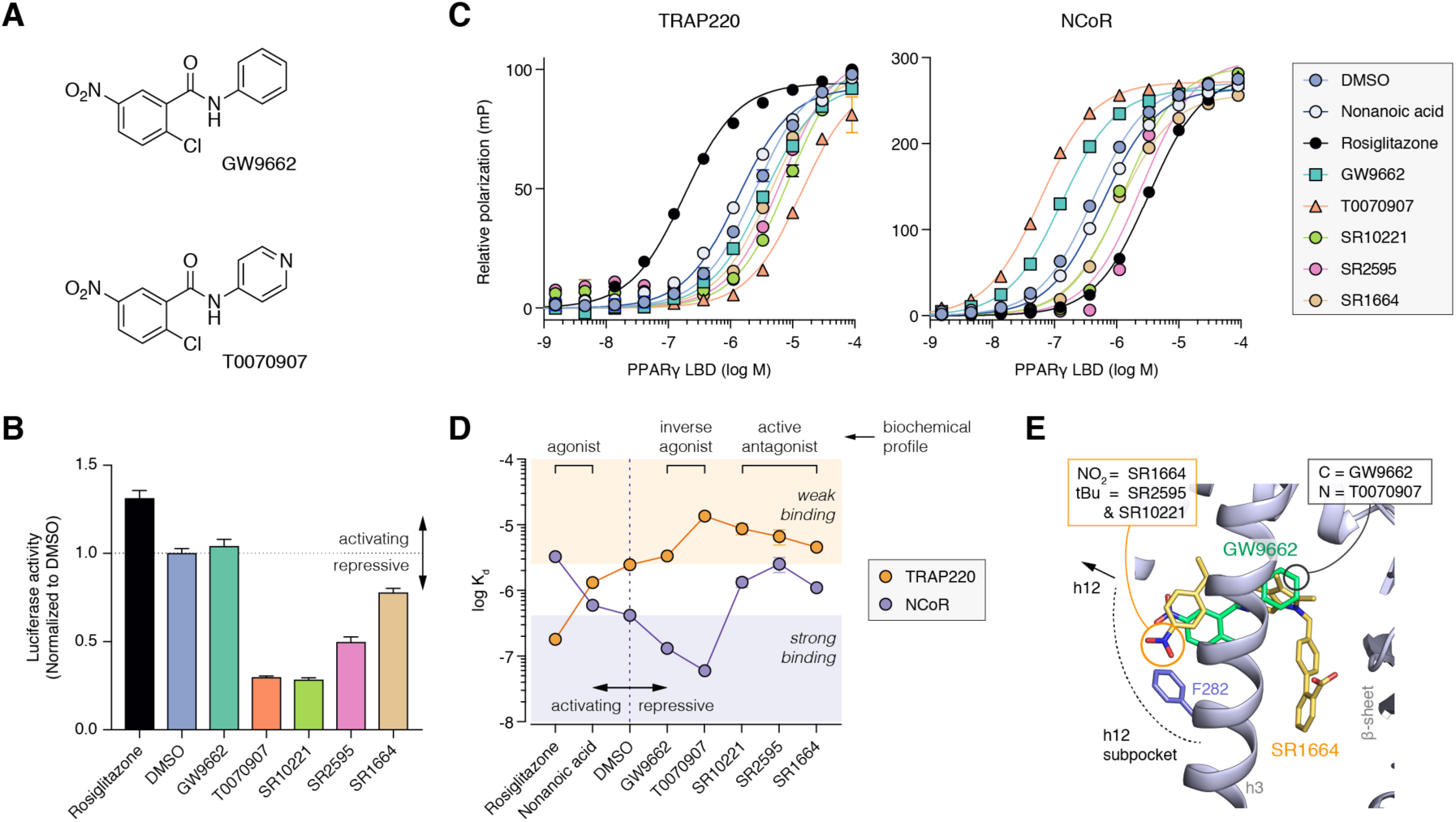
Differences in functional efficacy between GW9662, T0070907, and other synthetic PPARγ ligands. (**A**) Chemical structures of GW9662 and T0070907. (**B**) Cell-based luciferase transcriptional assay showing the effect of activating and repressive ligands (5 μM) on full-length PPARγ transcription in HEK293T cells. (**C**) Fluorescence polarization coregulator interaction assay showing the effect of ligands on the interaction between the PPARγ LBD and peptides derived from the TRAP220 coactivator or NCoR corepressor. (**D**) Kd values derived from the coregulator interaction assay. Dotted orange and purple lines note the DMSO/apo-PPARγ values. Orange and purple shaded areas note the affinity regions for an ideal inverse agonist (i.e., for weaker TRAP220 affinity, orange circles in the orange square; for higher NCoR affinity, purple circles in purple squares). (**E**) Superposition of crystal structures of the PPARγ LBD bound to GW9662 (PDB code 3B0R; ligand in green, cartoon in blue) and SR1664 (PDB code 4R2U; ligand in yellow). The location of the simple substitution between GW9662 (methine) vs. T0070907 (nitrogen) is marked with a black circle, and the SR2595 and SR10221 tert-butyl extension within the helix 12 subpocket towards F282 (blue) from the SR1664 parent compound is marked with a yellow circle.

We next characterized how the ligands affect the recruitment of peptides derived from the TRAP220 coactivator and the NCoR corepressor (Figure 1C), two coregulator proteins that influence PPARγ-mediated transcription ^20,21^. Compared to unliganded PPARγ (Figure 1D), nonanoic acid (a natural PPARγ agonist)^22^ and to a larger degree rosiglitazone (a synthetic PPARγ agonist) increased the affinity of TRAP220 and decreased the affinity of NCoR. Characteristic of an inverse agonist, T0070907 displayed an opposite profile to the agonists: decreasing the affinity of TRAP220 and increasing the affinity of NCoR. GW9662 displayed a similar trend as T0070907, though the changes in coregulator affinity were subtler: the affinity for NCoR increased, but unlike T0070907 the affinity for TRAP220 did not change (weaken) appreciably. Interestingly, the repressive compounds SR2595 and SR10221 have distinct profiles compared to T0070907: characteristic of a direct antagonist, they decreased affinity for both TRAP220 and NCoR.

We were interested in the structural mechanism by which the seemingly minor methine-to-nitrogen ligand substitution could switch a covalent neutral antagonist (GW9662) into a covalent inverse agonist (T0070907). Noncovalent analogs of the PPARγ agonist farglitazar ^16^ and the neutral antagonist/nonagonist SR1664 6 were designed to perturb the conformation of the helix 12/AF-2 surface. The farglitazar analogs contain ligand extensions towards helix 12. SR2595 and SR10221 contain a tert-butyl extension that perturbed the conformation of F282 on helix 3 (Figure 1E) located within the orthosteric pocket near the loop preceding helix 12, a region we refer to as the helix 12 subpocket. We previously showed that the F282/AF-2 steric clash caused by SR2595 and SR10221 binding increases the dynamics of helix 3, which is part of the AF-2 surface, thereby reducing the basal activity of PPARγ ^6^. Our coregulator recruitment data show this AF-2 clash affords a direct antagonist profile for SR2595 and SR10221 that leads to transcriptional repression (Figure 1B–D). However, the phenyl (GW9662) to pyridyl (T0070907) change is distant from F282, the helix 12 subpocket, or the AF-2 surface (Figure 1E), suggesting the inverse agonist profile of T0070907 may originate from a unique structural mechanism.

### A pyridyl-water hydrogen bond network unique to T0070907

To gain structural insight into the mechanism of action, we solved the crystal structure of T0070907 covalently bound to PPARγ to a resolution of 2.26 Å (PDB code 6C1I; Table S1) and compared our structure to an available crystal structure of GW9662-bound PPARγ (PDB code 3B0R). In both cases, PPARγ crystallized in the same space group and contained a dimer in the asymmetric unit with the expected alpha helical sandwich fold (Figure 2A). Structural superposition revealed nearly identical backbone conformations between the two structures (Cα r.m.s.d.: overall, 1.7 Å; chain A only, 1.34 Å; chain B only, 1.73 Å). In each of the structures, chain A adopts an “active” conformation with helix 12 docked into the activation function-2 (AF-2) surface, whereas helix 12 in chain B is distorted due to crystal packing interactions (Figure S2). Our crystals of T0070907-bound PPARγ were obtained by soaking the ligand into preformed apo-protein crystals since our cocrystallization attempts failed. Strong electron density was observed for T0070907 in chain B and lower, less defined electron density in chain A.

**Figure 2.**
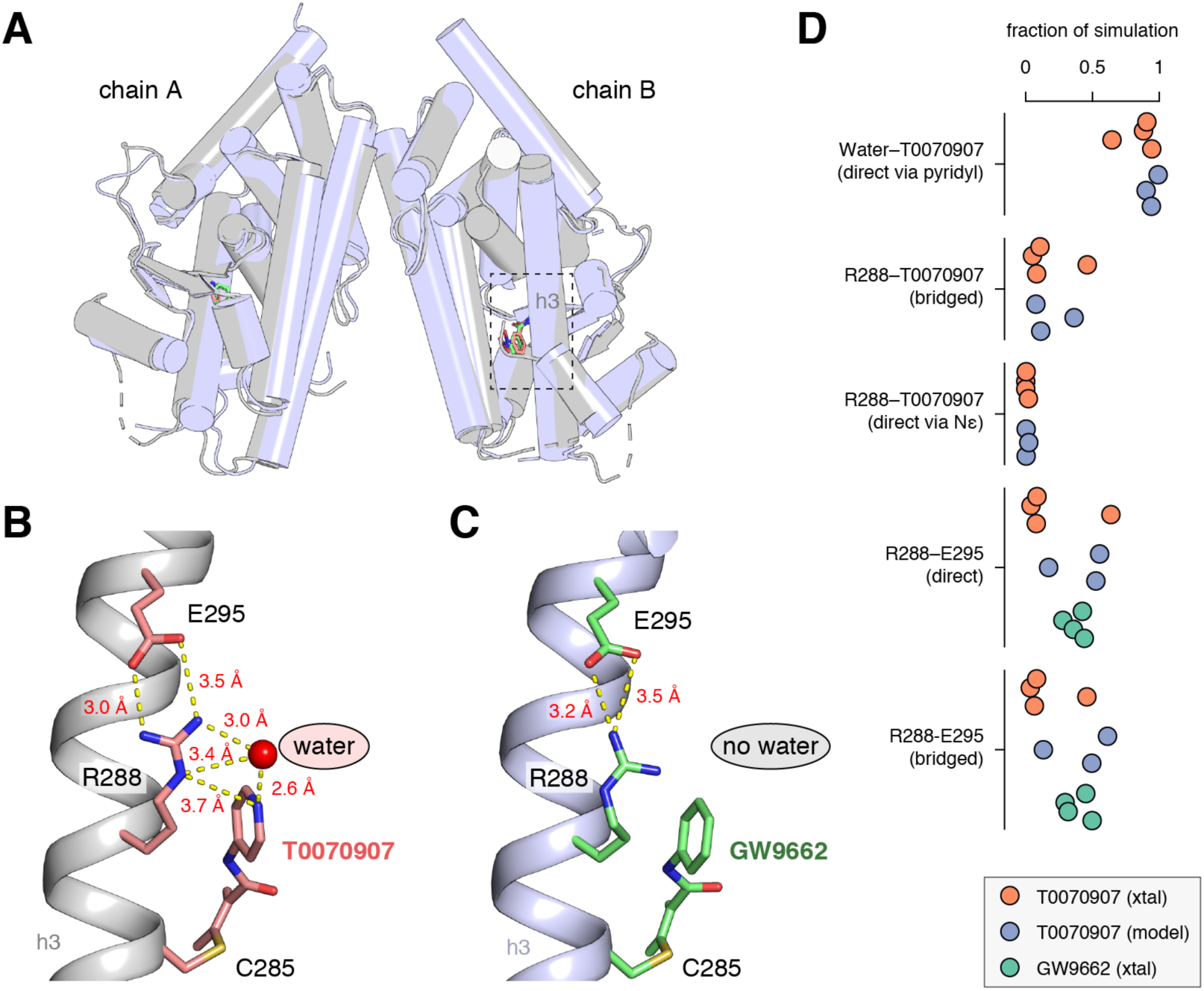
Crystal structure of T0070907-bound PPARγ LBD reveals a water-mediated pyridyl-protein hydrogen bond network. (**A**) Overall structure of T0070907-bound PPARγ (PDB code 6C1I; grey) and overlay with GW9662-bound crystal structure (PDB code 3B0R; blue). (**B**) A water-mediated hydrogen bond network in the T0070907-bound crystal structure (chain B is shown) links the pyridyl group in T0070907 to the R288 side chain, which forms a bipartite hydrogen bond with the E295 side chain. (**C**) The GW9662-bound crystal structure (chain B is shown) lacks the R288-ligand hydrogen bond network but contains the R288-E295 hydrogen bond. (**D**) Pyridyl-water network hydrogen bonds populated during molecular dynamics simulations for T0070907- and GW9662-bound structures starting from crystallized (xtal) and modeled (model) conformations.

Focusing on the pyridyl ring of T0070907, a water-mediated hydrogen bond network connects the pyridyl nitrogen to the Nε atom in the R288 side-chain (Figure 2B). Furthermore, the guanidinyl side chain of R288 forms a bipartite hydrogen bond with the side chain of E295. In contrast, in the GW9662-bound structure the hydrogen bond network is not extensive due the lack of a hydrogen bond acceptor in the phenyl ring of GW9662 (Figure 2C). To confirm the stability of the network observed in the crystal structures, we performed molecular dynamics simulations of T0070907- and GW9662-bound PPARγ ranging from 4–26 microseconds in length (Figure 2D) preserving the crystallized waters (xtal), as well as a model we generated of T0070907- bound PPARγ from the GW9662-bound PPARγ crystal structure solvated without crystallized waters (model). In the simulations, the pyridyl group of T0070907 was hydrogen bonded to a water molecule for a significant fraction of the simulation (65– 95%), as was the water-bridged R288-T0070907 pyridyl (5–46%). In contrast, a direct interaction between R288 and the pyridyl group of T0070907 not mediated by water was lowly populated (<2%). A direct (4–64%) and water-bridged (3–61%) R288-E295 interaction was also confirmed. The extensive pyridyl-based water-mediated hydrogen bond network is not possible to the hydrophobic phenyl group of GW9662, revealing a unique chemical feature in T0070907 that could confer inverse agonism.

### The pyridyl-water network is essential for inverse agonism

To test the functional role of the pyridyl-water hydrogen bond network observed in our T0070907-bound crystal structure, we generated variants of PPARγ by mutating residues that we predicted would maintain or break the pyridyl-water network. We hypothesized that mutation of R288 to a different positively charged residue (R288K) would maintain the pyridyl-water network, whereas mutation to a hydrophobic residue (R288A or R288L) would break the pyridyl-water network. If the hydrogen bond network is important for inverse agonism, we hypothesized that breaking this network via hydrophobic R288 mutations would afford a similar functional efficacy profile for both T0070907 and GW9662. We also generated a E295A mutation to test the importance of the bipartite hydrogen bond between R288 and E295. These mutations did not affect the structural integrity or stability of the PPARγ LBD as assessed by circular dichroism (CD) spectroscopy (Figure S3).

Using a cell-based transcription assay, we tested the combined effect of the mutants and covalent ligands on PPARγ cellular activation (Figure 3A). Wild-type PPARγ and the R288K mutant showed a similar profile, where cells treated with T0070907 showed decreased PPARγ transcription compared to DMSO or GW9662 treatment. In contrast, T0070907 did not decrease transcription for the R288A or R288L mutants, indicating the R288-mediated pyridyl-water network is responsible for the transcriptional repression conferred by T0070907. The E295A mutant maintained the cellular efficacy preference of T0070907 over GW9662, indicating that bipartite hydrogen bond is not a major contributor to stabilizing the repressive activity, though both GW9662 and T0070907 showed lower activity compared to wild-type PPARγ.

**Figure 3.**
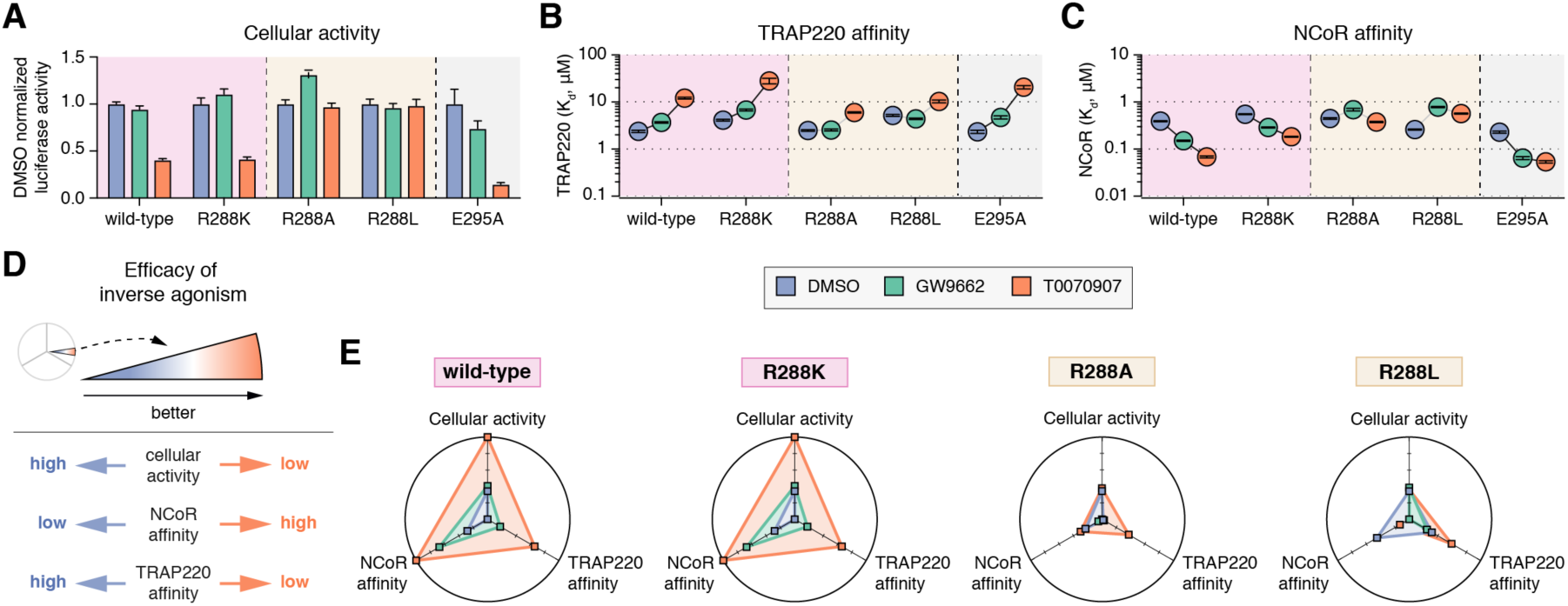
The R288-pyridyl interaction is essential for conferring inverse agonism. (**A**) HEK293T cells transfected with full-length PPARγ expression plasmid along with a 3x-PPRE-luciferase reporter plasmid and treated with the indicated ligands (5 μM). Data plotted as the average and s.e.m. of six experimental replicates. (**B,C**) Affinities determined from a fluorescence polarization assay of wild-type and mutant PPARγ LBDs preincubated with a covalent ligand (GW9662 or T0070907) or vehicle (DMSO) binding to FITC-labeled (**B**) TRAP220 or (**C**) NCoR. Data plotted as the Kd value and error from fitting data of two experimental replicates using a one site binding equation. (**D**) Legend to the “web of efficacy” radar chart diagrams. Inverse agonism is associated with data points populating the periphery of the plots. (**E**) Radar “web of efficacy” plots displaying assay data for wild-type PPARγ and mutant variants. Data normalized within the range of values for each assay (**A–C**).

We next tested the effect of the mutants on coregulator recruitment by determining binding affinities for the TRAP220 (Figure 3B) and NCoR (Figure 3C) peptides for wild-type PPARγ LBD and the mutant variants with or without pretreatment with GW9662 or T0070907. Consistent with the cell-based transcription assay, the R288K and E295A mutants maintained the inverse agonist coregulator binding profile of T0070907 and rank ordering. In contrast, the R288A and R288L mutants showed similar affinity for TRAP220 and NCoR when covalently bound to T0070907 or GW9662. This indicates the pyridyl-water network directs the inverse agonism profile of T0070907, the lack of which results in a neutral antagonist profile.

To more robustly compare how the wild-type and mutant PPARγ variants performed in the above assays, we performed a version of the “web of efficacy” analysis used in the G-protein coupled receptor (GPCR) field to study ligand signaling bias 23,24. We plotted the multivariate data on a radar chart with axes corresponding to each of the assays whereby conditions with the most efficacious (i.e., that are biased towards) inverse agonism properties populate the outer ring of the radar chart, and less favourable or efficacious conditions populate the center (Figure 3F). The analysis clearly shows that T0070907 selects more efficacious inverse agonism functions only for wild-type PPARγ and the R288K mutant variant (Figure 3G). This dramatic result reveals that R288-mediated pyridyl-water network directs inverse agonism conferred by T0070907.

### T0070907-bound PPARγ exchanges between two long-lived conformations

Despite the different inverse agonist and neutral antagonist profiles of T0070907 and GW9662, the pyridyl-water network is the primary structural difference observed in the crystal structures. In principle, there should be structural changes in the AF-2 coregulator interaction surface to account for the different pharmacological profiles. Notably, however, the conformation of the AF-2 surface, which includes helix 12, is influenced by crystal contacts (Figure S2). Furthermore, structural superposition of PPARγ crystal structures bound to pharmacologically distinct ligands shows no major structural changes that explain their different activities ^25^. However, NMR studies have shown that the orthosteric pocket and helix 12 are dynamic on the microsecond-to-millisecond (μs-ms) time scale in the ligand-free/apo-form, which results in very broad or unobserved NMR peaks for residues in helix 12 and within the orthosteric pocket and surrounding regions ^6,13^. Binding of a noncovalent full agonist stabilizes the helix 12/AF-2 surface resulting in the appearance of NMR peaks that were missing in the apo-form, whereas noncovalent partial agonists, neutral antagonists, and inverse agonists do not stabilize helix 12 ^6,13,25^.

We used NMR spectroscopy to assess the impact of T0070907 and GW9662 on the dynamics of the PPARγ LBD. NMR data of apo-PPARγ are similar to PPARγ covalently bound to GW9662 with widespread μs-ms dynamics in the ligand-binding pocket and helix 12/AF-2 surface ^26^. In remarkable contrast, T0070907-bound PPARγ showed a wide-spread stabilization of μs-ms dynamics, evident by the appearance of peaks in 2D [^1^H,^15^N]-transverse relaxation optimized spectroscopy-heteronuclear single quantum coherence (TROSY-HSQC) NMR spectra relative to GW9662-bound PPARγ (Figure 4A). This includes well-resolved T0070907-bound NMR peaks corresponding to residues located in close proximity to the T0070907 R288-mediated pyridyl-water network (Figure 4B) in the β-sheet (V248, G344, G346), helix 3 (I279, G284) and the adjacent helix 7 (G361). Furthermore, an NMR peak V322 on helix 5 within the AF-2 surface also appears, indicating stabilization of the AF-2 surface. The T0070907-bound crystal structure shows a larger network of water-mediated hydrogen bonds, which form a molecular hub linking the pyridyl-water network to the the β-sheet (via backbone hydrogen bonds to I341 and E343) and helix 5 (via backbone hydrogen bond to I326).

**Figure 4.**
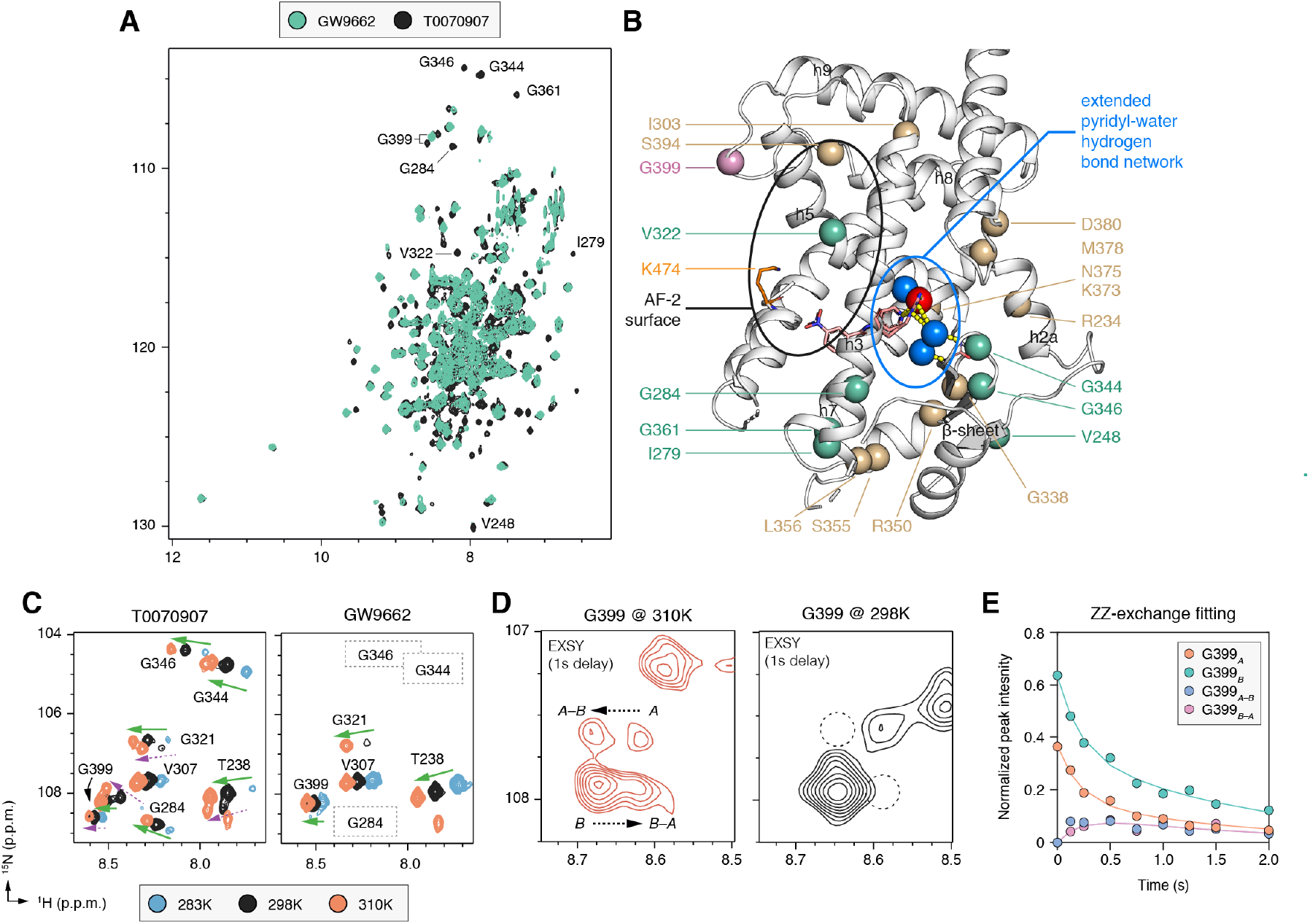
NMR detected exchange between two long-lived T0070907-bound conformations. (**A**) Overlay of [^1^H,^15^N]-TROSY-HSQC NMR spectra of ^15^N-PPARγ LBD bound to GW9662 or T0070907. (**B**) Binding of T0070907 but not GW9662 stabilizes intermediate exchange (μs-ms time scale) dynamics (residues labeled in (^A^) shown in green spheres) and causes peak doubling (tan and pink spheres; G399 is colored pink for emphasis). Data plotted on the T0070907-bound PPARγ crystal structure and important structural regions are highlighted as follows: AF-2 surface (black oval); an extended pyridyl-water hydrogen bond network (blue spheres, yellow dotted lines, blue oval), beyond the key pyridyl-water interaction (red sphere). (^C^) Snapshot overlays of [^1^H,^15^N]-TROSY-HSQC spectra of 15N-PPARγ LBD bound to T0070907 or GW9662. The spectral region displayed shows single peaks are observed whenbound to either ligand (denoted with green arrows), temperature-dependent NMR peak doubling when bound to T0070907 (purple dotted arrows), and absent peaks due tointermediate exchange on the NMR time scale when bound to GW9662 (dotted rectangles) (^D^) Snapshots of ZZ-exchange ^15^N-HSQC NMR spectra (delay = 1 s) of T0070907-bound ^15^N-PPARγ LBD focused on G399 at the indicated temperatures. Two G399 conformational statesare denoted as *A* and *B* with the ZZ-exchange transfer crosspeaks as *A–B* and *B–A*. (*E*) ZZ-exchange NMR analysis build-up curve from for G399 at 310K generated by plotting peak intensities of the state*A*and*B*peaks and exchange crosspeaks (*A–B*and*B-A*) as a function of delay time.

Temperature-dependent NMR studies (Figure 4C) further revealed a number of PPARγ residues experience peak doubling when bound to T0070907, but not when bound to GW9662, including G399 located near the AF-2 surface (Figure 4B). Peak doubling indicates the presence of two long-lived T0070907-bound structural conformations in slow exchange on the NMR time scale, where the difference in chemical shift between the two states (*Δv*, in Hz) is much greater than the exchange rate (*k_ex_*) between conformations on the order of milliseconds-to-seconds (ms–s) ^27^. Thus, the increase in NMR peaks when PPARγ is bound to T0070907 is not only due to stabilization of μs-ms dynamics but also the presence of two long-lived structural conformations.

ZZ-exchange NMR experiments (also called EXSY, or exchange spectroscopy) enable detection of the interconversion between long-lived structural states via transfer of the 1H chemical shift of one state to the other when *k_ex_*» 0.2−100 s-1 and k_ex_>>*Δv*. Exchange crosspeaks for G399, which shows well dispersed peak doubling, were observed at 37°C but not at 25°C (Figure 4D), indicating the exchange between the two conformations is too slow to be measured at room temperature (*k_ex_*< 0.2/s). To determine an exchange rate, we performed ZZ-exchange experiments with varying exchange delays at 37°C and fit the data to a two-state interconversion model (Figure 4E), which provided an exchange rate of ~2.1/s between the upfield shifted state (*P_A_* = 37%; *k*_*A*→*B*_ = 0.8/s) and downfield shifted state (*P_B_* = 63%; *k*_*B*→*A*_ = 1.3/s). Notable peak doubling, many of which show well dispersed exchange crosspeaks, is widespread through the PPARγ LBD (Figure S4A), though in most cases spectral overlap did not permit fitting of the data to extract an exchange rate. However, residues with notable peak doubling comprise distant structural regions that are also connected via the aforementioned extended pyridyl-water network (Figure 4B), including the β-sheet (G338) and helix 6 (R350, S355, L356) within the ligand-binding pocket; a surface comprising helix 2a (R234) and the C-terminal region of helix 7 and the loop connecting helix 7 and 8 (K373, N375, E378, D380); helix 3 near the AF-2 surface (I303); and the loop connecting helix 8 and 9 near the AF-2 surface (S394), which also includes G399. In total, the NMR analysis revealed that T0070907-bound PPARγ undergoes a global conformational change between two long-lived structural conformations.

### T0070907 populates a mutual conformation with GW9662 and a unique conformation

G399 is an ideal NMR observable probe that is sensitive to the conformation of the AF-2 surface: it is structurally proximal and linked to the AF-2 surface through water-mediated hydrogen bonds to N312 and D311 on helix 5, but does not directly interact with a bound coregulator peptide (Figure 5A). Strikingly, for G399 and the other residues that showed peak doubling in the ZZ exchange analysis, we found that the backbone amide chemical shifts of one of the two peaks observed for T0070907-bound PPARγ are similar to the single peak observed for GW9662-bound PPARγ (Figure 5B and Figure S4B). This indicates that one of the long-lived T0070907-bound conformations is structurally similar to GW9662-bound PPARγ, which below we refer to as the mutual conformation, and the other conformation is uniquely populated only when bound to T0070907.

We also assessed the conformational state of helix 12 directly using ^19^F NMR (Figure 5C) by attaching the ^19^F NMR-detectible probe 3-bromo-1,1,1-trifluoroacetone (BTFA) on K474 (Figure 5A). The ^19^F spectral profile of GW9662-bound PPARγ revealed two peaks corresponding to a major state (right peak; 78%) and minor state (left peak; 22%). T0070907-bound PPARγ also shows two peaks with chemical shift values similar to GW9662-bound PPARγ, but the population magnitudes of the states are switched and skewed towards the left peak. Strikingly, this helix 12/AF-2 surface ^19^F NMR probe showed the same relative population sizes observed in the G399/proxy to the AF-2 surface ZZ-exchange analysis (34% and 66%, respectively). The right peak abundantly populated by GW9662 and moderately populated by T0070907 likely corresponds to the mutual G399 conformation from the 2D NMR analysis. In contrast, the left ^19^F NMR peak likely corresponds to the unique G399 conformation; this peak is abundantly populated by T0070907 but lowly populated by GW9662. The low abundance of this peak when bound to GW9662 could explain in part why it was not detected by the 2D NMR analysis, which has lower overall sensitivity of signal-to-noise compared to the ^19^F NMR analysis. However, the BTFA probe attached to helix 12 may also be sensitive to larger structural changes, and thus chemical environments, compared to backbone amide of G399.

**Figure 5.**
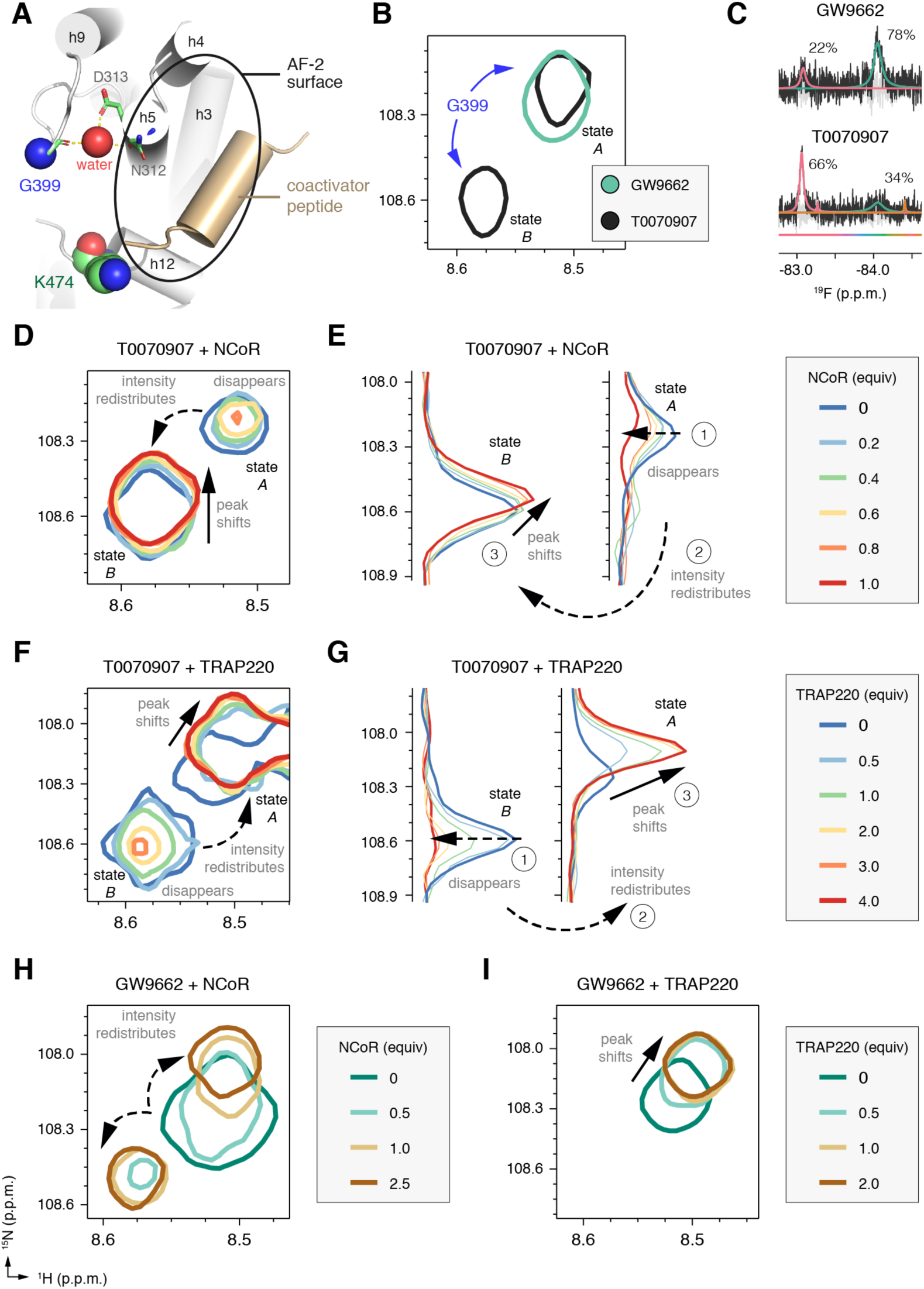
T0070907 but not GW9662 prepopulates a corepressor-bound conformation. (**A**) Structural location of G399, which is connected to the AF-2 coregulator interaction surface via water-mediated hydrogen bonds to N312 and D313 but does not directly interact with a coregulator peptide bound to the PPARγ LBD (PDB code 2PRG). (**B**) Snapshot overlay of [^1^H,^15^N]-TROSY-HSQC NMR spectra of ^15^N-PPARγ LBD bound to GW9662 or T0070907 shows that the single GW9662-bound G399 peak has similar chemical shift values to one of the two T0070907-bound G399 peaks (state *A*) whereas state *B* is uniquely populated by T0070907. (**C**) Deconvoluated ^19^F NMR data of BTFA-labeled PPARγ LBD covalently bound to GW9662 or T0070907. (**D–I**) Snapshots of [^1^H,^15^N]-TROSY-HSQC spectra of ^15^N-PPARγ LBD bound to (**D,E**) T0070907 and titrated with NCoR peptide; (**F,G**) T0070907 and titrated with TRAP220 peptide; (**H**) GW9662 and titrated with NCoR peptide; and (**I**) GW9662 and titrated with TRAP220 peptide. 1D spectra in (**E**) and (**G**) show ^15^N planes extracted from (**D**) and (**F**), respectively, to better illustrate the peak transitions: **1**, disappearance of the state *A* or *B* peak; **2**, the redistribution of their intensities to the other state; and **3**, the slight shifting of the other state towards, increased population of, the peptide-bound state.

### The unique T0070907 conformation prepopulates a corepressor-bound conformation

We wondered whether the unique and mutual long-lived T0070907-bound conformations would display similar or distinct coregulator interaction preferences. To test this, we titrated the NCoR corepressor and TRAP220 coactivator peptides and monitored their binding to ^15^N-PPARγ LBD by NMR. Remarkably, titration of NCoR corepressor peptide into T0070907-bound PPARγ (Figure 5D,E) resulted first in a shifting of the unique G399 conformation (peak *B*) towards a similar chemical shifts values and intensity, nearly saturating around 0.6 equiv NCoR, which is the approximate population (*P_B_*) of the state from the ZZ-exchange analysis. Only minor changes in peak intensity were observed for the mutual peak (peak *A*) until the titration reached 0.6 equiv NCoR peptide, at which point this peak decreased in intensity with a concomitant increase in the intensity of the unique conformation peak that shifted to the NCoR-bound conformation. This second transition (peptide free state *A*→ NCoR bound state) saturated at 1 equiv of NCoR peptide in slow exchange on the NMR time scale. These results are consistent with our coregulator affinity data showing mid-nM affinity for NCoR binding.

We next examined the effect of TRAP220 peptide binding to T0070907-bound PPARγ (Figure 5F,G). In contrast to the NCoR results, the mutual conformation of G399 (state *A*) transitioned first to a peak with similar chemical shift values and intensity. This first transition saturated at the first titration point (0.5 equiv), which is slightly larger than the approximate population (*P_A_*) of the state from the ZZ-exchange analysis. A second transition also occurred where the unique conformation (state *B*) showed a decrease in peak intensity, but only at TRAP220 amounts more than 0.5 equiv, which resulted in a concomitant increase in the intensity of the mutual peak that shifted to the TRAP220-bound peak. This second transition did not saturate until 4 equiv of TRAP220 was added, indicating that the unique conformation displays much weaker affinity for TRAP220 relative to the mutual conformation. Notably, the same coregulator binding trends for the other residues with peak doubling, where NCoR binding shifts the peak populations towards the unique state *B* (Figure S4C) and TRAP220 binding shifts the peak populations towards the mutual state *A* (Figure S4D).

Finally, we examined the binding of GW9662-bound PPARγ with NCoR (Figure 5H) and TRAP220 (Figure 5I). Interestingly, whereas NCoR or TRAP200 binding to T0070907-bound PPARγ consolidated the unique and mutual conformations into one peptide-bound conformation, NCoR binding to GW9662-bound PPARγ caused peak doubling of the single GW9662-bound G399 NMR peak towards chemical shift values similar to the NCoR- and TRAP220-bound form of T0070907-bound PPARγ. In contrast, TRAP220 binding only shifted the single GW9662-bound G399 peak towards the TRAP220-bound form of T0070907-bound PPARγ.

These dramatic results reveal that the two long-lived T0070907-bound conformations have different binding preferences for NCoR and TRAP220. The NMR chemical shit perturbation profiles reveal that the unique conformation has strong affinity for NCoR but weak affinity for TRAP220, whereas the mutual conformation has moderate affinity for NCoR and strong affinity for TRAP220. Moreover, the NMR chemical shifts of the unique and mutual T0070907-bound conformations in the absence of coregulator peptide are similar to the NCoR- and TRAP220-bound forms, respectively. This indicates that the unique and mutual T0070907-bound states prepopulate a corepressor-like and coactivator-like bound conformation that individually afford privileged high-affinity binding to NCoR and TRAP220, respectively. Furthermore, the chemical shift difference between the unique T0070907 conformation and NCoR-bound state (i.e., the degree of state *B* shifting) is much smaller than the mutual conformation and TRAP220-bound state (i.e., the degree of state *A* shifting). This indicates the corepressor-like conformation prepopulated by T0070907 is more similar to the corepressor-bound state than the coactivator-like conformation prepopulated by T0070907 is to the coactivator-bound state. In contrast, NCoR binding to GW9662-bound PPARγ introduces a “conformational frustration” within the AF-2 surface. In addition to not prepopulating the corepressor-bound conformation, the AF-2 surface of GW9662-bound PPARγ is found in both the corepressor- and coactivator-bound conformations upon binding NCoR—both of which could contribute to the neutral antagonism profile for GW9662 as opposed to the enhanced inverse agonism profile of T0070907 derived through prepopulation of conformational states that have high affinity for corepressor and low affinity for coactivator.

## Discussion

Carbon (methine)-to-nitrogen ligand substitutions are known to have beneficial effects on pharmacological parameters^28^, though it is difficult to predict how subtle changes in chemical structure impact switches in functional efficacy^29,30^. GW9662 and T0070907 are widely used as chemical tools due to their ability to covalently bind to PPARγ and inhibit ligand binding to the orthosteric ligand-binding pocket. Although it is generally acknowledged that these highly similar compounds display distinct PPARγ activity profiles, the molecular basis for their functional differences in efficacy has remained elusive. Our studies, which combine crystallography, molecular dynamics simulations, NMR spectroscopy, and mutagenesis coupled with biochemical and cellular assays, illuminate a novel structural mechanism affording the inverse agonism of T0070907. Our crystal structure of T0070907-bound PPARγ revealed a water-mediated hydrogen bond network linking the critical inverse agonist “switch” residue, R288, which is distal from the activation function-2 (AF-2) coregulator interaction surface, to the pyridyl group of T0070907. Our NMR analysis shows that T0070907-bound PPARγ, but not GW9662-bound PPARγ, slowly exchanges between two long-lived conformations. One of these conformations is shared with GW9662-bound PPARγ and similar to the coactivator-bound state. The other conformation is uniquely and abundantly populated by T0070907 and highly similar to the corepressor-bound state, thus affording higher affinity corepressor binding and inverse agonism for T0070907.

We demonstrated the importance of the pyridyl-water network in directing inverse agonism of PPARγ using mutagenesis coupled with functional assays. However, no major structural differences were observed in the crystal structures that explain their functional profiles. In contrast, our NMR data revealed clearly that T0070907-bound PPARγ, but not GW9662-bound PPARγ, exchanges between two long-lived conformations, one of which prepopulates a conformation similar to the corepressor-bound state. The active AF-2 conformation bound to a coactivator peptide has captured in numerous crystal structures of PPARγ and other nuclear receptors. Thus far, no structures have been reported for PPARγ bound to a corepressor peptide and, relative to coactivator-bound structures, a limited number of structures have been reported for corepressor-bound nuclear receptors ^31^. However, from these studies it is known that the binding regions of coactivator and corepressor (NCoR and SMRT) peptides overlap. In the corepressor-bound structures, helix 12 is displaced from its “active” conformation and shows variability in its crystallized conformation, which may reflect a dynamic “inactive” conformational profile. Our NMR studies show that as a whole the AF-2 surface, using G399 as a proxy, is primed for high affinity binding to NCoR when PPARγ is bound to T0070907.

There are two main underlying conclusions from our NMR findings on T0070907-bound PPARγ. First, the crystallized AF-2 conformations do not likely represent the conformations of T0070907-bound PPARγ in solution, since crystal packing influence the position of helix 12. Our differential NMR analysis not only revealed peak doubling for T0070907-bound PPARγ but also a stabilization of μs-ms timescale dynamics relative to GW9662-bound PPARγ. Furthermore, using ^19^F NMR we showed that helix 12 dynamically exchanges between two long-lived conformations, one of which is significantly populated only by T0070907. Second, we used molecular dynamics simulations to verify the structural integrity of the pyridyl-water network. However, given that the NMR-detected exchange rate between the two T0070907-bound conformations is >1s, access to helix 12/AF-2 conformations that would be consistent with our NMR data in molecular simulations is inaccessible with current standard simulation approaches. Overall, our work shows that the combination of different but complementary structural methods—crystallography, NMR, and molecular simulations—provides the full picture of ligand mechanism of action, which as we demonstrate here, involves a simultaneous prepopulation of long-lived conformational states with distinct functions.

Our findings suggest a new means for pharmacologically directing transcriptional repression via inverse agonism of PPARγ. The previous finding that ligand engagement of, or hydrogen binding to, helix 12 via Y473 is critical for mediating agonism transformed the way that PPARγ agonists were developed ^32^. The AF-2 steric clash mechanism of action for the repressive PPARγ compounds SR2595 and SR10221 ^6,16^ shows a coregulator interaction profile consistent with a direct antagonist rather than an inverse agonist. In contrast, our studies here indicate that ligand hydrogen bonding to the guanidinyl side chain of R288, water-mediated or perhaps directly, may be a critical mediator of inverse agonism. Repressive PPARγ modulators show promise for improving the therapeutic index associated with anti-diabetic PPARγ ligands by promoting bone formation rather than decreasing bone mass ^5,6^, which occurs with agonists used clinically such as the TZDs. Furthermore, repression of PPARγ activity affects fat mobilization and may be a means to therapeutically treat obesity and extend lifespan ^7^, and T0070907 has demonstrated efficacy in cancer models ^9-11^. Thus, our findings should inspire future work to develop and characterize inverse agonists to probe the repressive functions of PPARγ.

## Methods

### Materials and reagents

Human PPARγ LBD (residues 203–477 in isoform 1 numbering, which is commonly used in published structural studies and thus throughout this manuscript; or residues 231–505 in isoform 2 numbering) or mutant proteins were expressed in *Escherichia coli* BL21(DE3) cells as TEV-cleavable hexahistidine-tagged fusion protein using a pET46 plasmid as previously described ^13,26^. The final storage buffer for samples following size exclusion chromatography and subsequently frozen at −80 °C was 50 mM potassium chloride (pH 7.4), 20 mM potassium phosphate, 5 mM TCEP, and 0.5 mM EDTA. Covalent ligand treatment was performed overnight at 4 °C with a 2X molar excess of compound dissolved in d_6_-DMSO. Mammalian expression plasmids included Gal4-PPARγ-hinge-LBD (residues 185-477 in isoform 1 numbering; 213-505 in isoform 2 numbering) inserted in pBIND plasmid; and full-length PPARγ (residues 1-505; isoform 2) inserted in pCMV6-XL4 plasmid. Mutant proteins were generated using site directed mutagenesis of the aforementioned plasmids. GW9662, T0070907, and SR1664 were obtained from Cayman Chemical; rosiglitazone was obtained from Tocris Bioscience and Cayman Chemical; SR2595 and SR10221 were previously synthesized in house ^6^.

Peptides of LXXLL-containing motifs from TRAP220 (residues 638–656; NTKNHPMLMNLLKDNPAQD) and NCoR (2256–2278; DPASNLGLEDIIRKALMGSFDDK) containing a N-terminal FITC label with a six-carbon linker (Ahx) and an amidated C-terminus for stability were synthesized by LifeTein.

### Cell-based transcriptional luciferase reporter assay

HEK293T cells were cultured in DMEM medium supplemented with 10% fetal bovine serum (FBS) and 50 units ml^−1^ of penicillin, streptomycin, and glutamine. Cells were grown to 90 % confluency and then seeded in 10 cm dishes at 4 million cells per well. Cells were transfected using X-tremegene 9 (Roche) and Opti-MEM (Gibco) with pCMV6 full-length PPARγ expression plasmid (4.5 μg) and 3xPPRE-lucifease reporter pGL2 plasmid (4.5 μg) and incubated for 18 h; plasmids were obtained from P. Griffin (Scripps) as used in previous studies. ^3,6,13,26^ Cells were transferred to white 384-well plates (Thermo Fisher Scientific) at 10,000 cells/well in 20 μL and incubated for 4 hr. Ligand (5 μM) or vehicle control was added (20 μL), cells incubated for 18 hr and harvested for luciferase activity quantified using Britelite Plus (Perkin Elmer; 20 μL) on a BioTek Synergy Neo multimode plate reader (Biotek). Data were analyzed using GraphPad Prism (luciferase activity vs. ligand concentration) and fit to a sigmoidal dose response curve.

### Fluorescence polarization (FP) coregulator interaction assay

The assay was performed in black 384-well plates (Greiner) in assay buffer (see above). His-PPARγ LBD was pre-incubated with or without a 2X molar excess of covalent ligand overnight at 4 °C and diluted by serial dilution. Noncovalent compounds were incubated with a constant concentration of 90 μM, equivalent to the maximum protein concentration, to ensure full occupancy. FITC-labeled NCoR and TRAP220 peptides were plated at a final concentration of 100 nM. Plates were incubated for 2 hr at 4 °C and measured on a BioTek Synergy Neo multimode plate reader at 485 nm emission and 528 nm excitation wavelengths. Data were plotted using GraphPad Prism and fit to one-site binding equation.

### Crystallography

PPARγ LBD protein was concentrated to 10 mg/ml and buffer exchange into phosphate buffer (20 mM KH_2_PO_4_/K_2_HPO_4_, pH 8, 50 mM KCl, 5 mM TCEP). Apo-PPARγ crystals were obtained after 3–5 days at 22°C by sitting-drop vapor diffusion against 50 μl of well solution using 96-well format crystallization plates. The crystallization drops contain 1 μl of protein sample mixed with 1 μl of reservoir solution containing 0.1 M MOPS, 0.8 M sodium citrate at pH 6.5. T0070907 was soaked into the PPARγ apo-crystals drop by adding 1 μl of compound at a concentration of 1 mM suspended in the same reservoir solution for 3 weeks. Crystals were cryoprotected by immersion in mother liquor containing 12% glycerol and flash-cooled in liquid nitrogen before data collection. Data collection was carried out at Beamlines 5.0.1 of BCSB at the Advanced Light Source (Lawrence Berkeley National Laboratory). Data were processed, integrated, and scaled with the programs Mosflm and Scala in CCP4 ^34,35^. The structure was solved by molecular replacement using the program Phaser ^36^ implemented in the PHENIX package ^37^ and used previously published PPARγ LBD structure (PDB code: 1PRG) ^38^ as the search model. The structure was refined using PHENIX with several cycles of interactive model rebuilding in COOT ^39^.

### Molecular dynamics simulations

A crystal structure of GW9662-bound PPARγ (PDB code 3B0R) along with our crystal structure T0070907 bound to PPARγ (PDB code 6C1I) were used to build initial structures in all simulations in this study. Two models were generated using 3B0R crystal structure. In the first model, chain A of 3B0R was used and GW9662 was transformed to T0070907 by converting phenyl ring of GW9662 to the pyridine ring of T0070907. The second 3B0R generated model was built using chain B conformation. In addition, chain B of the T0070907 crystal structure was used for a third build. The crystalized water molecules were kept in the models in which chain B conformations were used. The Modeller ^40^ extension within UCSF Chimera ^41^ was used to fill in the missing part of the protein in PDB files. The resulting structures were submitted to H++ server ^42^ to determine the protonation states of titratable residues at pH 7.4. AMBER names were assigned to different protonation states of histidine using pdb4amber provided in AmberTools 14 ^3^). In order to parametrize T0070907 and GW9662, the C285 with covalently attached ligand was protonated and methyl caps were added, saved as a separate PDB file using Chimera, and submitted to the R.E.D server ^43^ to calculate RESP ^44^ charges. AMBER cysteine residue values were used for the RESP charges for the cysteine backbone. The output mol2 file was used to generate the ac and prepin files following a method in the tutorial (http://ambermd.org/tutorials/basic/tutorial5/). Parmchk2 was used to create two force modification files from the prepin file, one that used AMBER ff14SB ^45^ parameter database values and another that used general Amber force field ^46^ (GAFF2) values, then Tleap was used to generate topology and coordinate files. The ff14SB force field was used to describe the protein. The resulting structure was solvated in a truncated octahedral box of TIP3P water molecules with the 10 Å spacing between the protein and the boundary, neutralized with Na+ and K+ and Cl− ions were added to 50 mM. The system was minimized and equilibrated in nine steps at 310 K with nonbonded cutoff of 8 Å. In the first step the heavy protein atoms were restrained by a spring constant of 5 kcal/mol Å2 for 2000 steps, followed by 15 ps simulation under NVT conditions with shake, then two rounds of 2000 cycles of steepest descent minimization with 2 and 0.1 kcal/mol Å^2^ restraints were performed. After one round without restraints, three rounds of simulations with shake were conducted for 5 ps, 10 ps and 10 ps under NPT conditions and restraints of 1, 0.5 and 0.5 kcal/mol Å^2^ on heavy atoms. Finally, an unrestrained NPT simulation was performed for 200 ps. Production runs were carried out with hydrogen mass repartitioned ^47^ parameter files to enable 4 fs time steps. Constant pressure replicate production runs were carried out with independent randomized starting velocities. Pressure was controlled with a Monte Carlo barostat and a pressure relaxation time (taup) of 2 ps. Temperature was kept constant at 310 K with Langevin dynamics utilizing a collision frequency (gamma_ln) of 3 ps-1. The particle mesh ewald method was used to calculate non-bonded atom interactions with a cutoff (cut) of 8.0 Å. SHAKE ^48^ was used to allow longer time steps in addition to hydrogen mass repartitioning. Analysis of trajectories was performed using cpptraj ^49^. Hydrogen bond analysis was performed using dist = 3.5 Å and angle = 100°^50^.

### CD Spectroscopy

Protein samples pre-incubated with or without a 2X molar excess of covalent ligand overnight at 4 °C were diluted to 10 μM in CD buffer (20 mM KPO4 pH 7.4, 10 mM KCl, 1 mM TCEP, 10% glycerol) and measured on a JASCO J-815 CD spectrometer by scanning from 190 nm to 300 nm at 20 °C or by increasing the temperature from 20 to 80 °C at 1 °C/min while monitoring the CD signal at 223 nm. Protein unfolding/melting temperature (Tm) was determined by fitting the data to a thermal unfolding equation ^51^ in GraphPad Prism.

### NMR spectroscopy

2D [^1^H,^15^N]-TROSY HSQC NMR data of 200 μM ^15^N-labeled PPARγ LBD, pre-incubated with a 2X molar excess of covalent ligand overnight at 4 °C, were acquired at 298K (unless otherwise indicated) on a Bruker 700 MHz NMR instrument equipped with a QCI cryoprobe in NMR buffer (50 mM potassium phosphate, 20 mM potassium chloride, 1 mM TCEP, pH 7.4, 10% D_2_O). For peptide titrations, peptides were dissolved in NMR buffer. ZZ-exchange experiments were acquired at 298K or 310K on Bruker 700 or 800 MHz NMR instrument equipped with a QCI or TCI cryoprobe, respectively, using exchange mixing times ranging from 0–2 s. Data were processed and analyzed using Topspin 3.0 (Bruker Biospin) and NMRViewJ (OneMoon Scientific, Inc.) ^52^, respectively. NMR chemical shift assignments previously described for ligand-bound PPARγ ^6,13,26^ were assigned to the spectra for well resolved residues with consistent NMR peak positions the presence of different ligands using the minimum chemical shift perturbation procedure williamson^53^. ZZ exchange data were fit to an exchange model for slow two-state interconversion ^27,54^ using a protocol described by, and a MATLAB script provided by, Gustafson, et al. ^55^.

For ^19^F NMR, PPARγ LBD K474C mutant protein was used to allow covalent attachment of 3-bromo-1,1,1-trifluoroacetone (BTFA) helix 12 via K474C. Mass spectrometry verified that GW9662 and T0070907 (2X molar excess) do not covalently attach to K474C (using a K474C/C285S double mutant protein that is in capable of covalent attachment to C285); using wild-type protein confirmed covalently attachment to C285. Samples were first incubated with 2X GW9662 or T0070907, then incubated with 2X BTFA, followed by buffer exchange into ^19^F NMR buffer (25 mM MOPS, 25 mM KCl, 1 mM EDTA, pH 7.4, 10% D2O). 1D ^19^F NMR data of 150 μM BTFA-labeled PPARγ LBD bound to GW9662 or T0070907 were acquired at 298K on a Bruker 700 MHz NMR instrument equipped with QCI-F cryoprobe. Chemical shifts were calibrated using an internal separated KF reference in ^19^F NMR buffer without TCEP contained in a coaxial tube inserted into the NMR sample tube. KF was set to be ––119.522 ppm, which is the chemical shift of the KF signal with respect to the ^19^F basic transmitter frequency for instrument. 1D ^19^F spectra were acquired utilizing the zgfhigqn.2 pulse program provided in Topspin 3.5 (Bruker Biospin). Data were processed using Topspin and deconvoluted with decon1d ^56^.

## Acknowledgements

We thank Sarah Mosure and Paola Munoz-Tello for critical reading of the manuscript. This work was supported by National Institutes of Health (NIH) grants R01DK101871 (DJK), F32DK108442 (RB), and R00DK103116 (TH); American Heart Association (AHA) fellowship award 16POST27780018 (RB); and the William R. Kenan, Jr. Charitable Trust (TSRI High School Student Summer Internship Program). A portion of this work (ZZ exchange) was performed at the National High Magnetic Field Laboratory (NHMFL/MagLab), which is supported by National Science Foundation (NSF) Cooperative Agreement No. DMR-1157490 and the State of Florida. ^19^F NMR data presented herein were collected at the CUNY ASRC Biomolecular NMR Facility.

## Author contributions

R.B. performed the cellular and biochemical assays. J.S. and J.F. performed crystallography. J.S. and D.J.K. performed the 2D NMR experiments. I.M.C. and T.S.H. performed the 19F NMR experiments. Z.H., T.S.H, and D.J.K. performed the molecular dynamics simulations. R.B., J.S., J.F., J.B., A.C., and I.M.C. performed mutagenesis and/or purified proteins. A.-L.B., P.R.G., and T.M.K. supplied compounds. R.B. and D.K. conceived the experiments and wrote the manuscript with input from all authors.

---

**Table S1.** X-ray data collection and refinement statistics.

|  |  |
| --- | --- |
| | T0070907-PPARγ LBD |
| **Data collection** | ALS-BCSB 5.0.1 |
| Space group | C 1 2 1 |
| Cell dimensions |  |
| $a, b, c$ (Å) | 92.99, 61.69, 118.66 |
| $\alpha, \beta, \gamma$ (°) | 90, 102.31, 90 |
| Resolution | 45.43–2.26 (2.341–2.26) |
| $R_{\text{pim}}$ | 0.02979 (0.3361) |
| I / $\sigma$(I) | 10.73 (2.12) |
| CC1/2 in highest shell | 0.906 |
| Completeness (%) | 99.53 (99.84) |
| Redundancy | 2.0 (2.0) |
| **Refinement** |  |
| Resolution (Å) | 2.26 |
| No. of reflections | 61110 |
| $R_{\text{work}}/R_{\text{free}}$ (%) | 21.54/28.38 |
| No. of atoms |  |
| Protein | 4181 |
| Water | 308 |
| $B$-factors | |
| Protein | 32.35 |
| Ligand | 44.64 |
| Water | 30.64 |
| Root mean square |  |
| Bond lengths (Å) | 0.008 |
| Bond angles (°) | 0.94 |
| Ramachandran favored | 95.11 |
| Ramachandran outliers | 0.59 |
| PDB accession code | 6C1I |
| *Values in parentheses indicate highest resolution shell.* |  |

**Figure S1.**
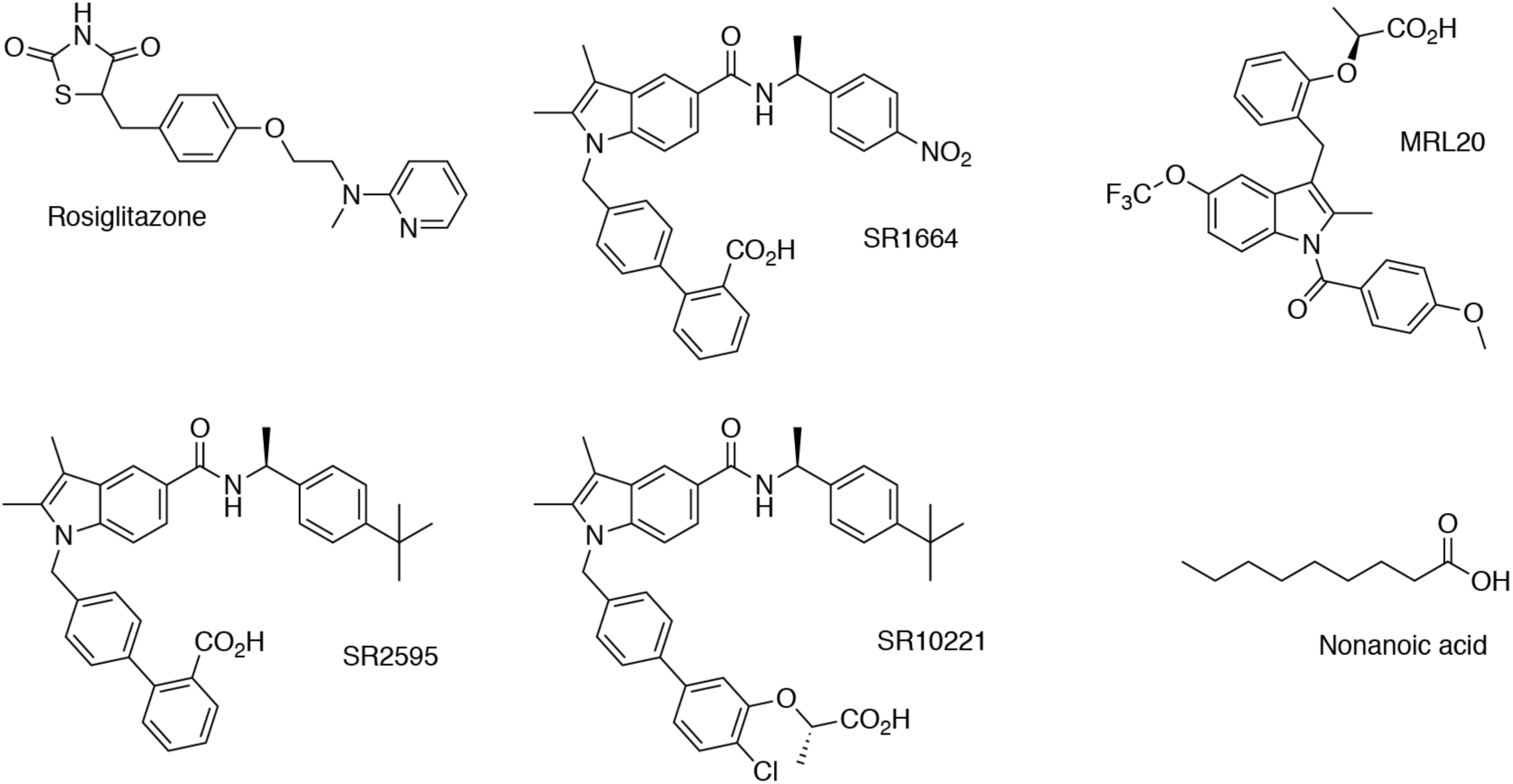
Noncovalent ligands used in the study.

**Figure S2.**
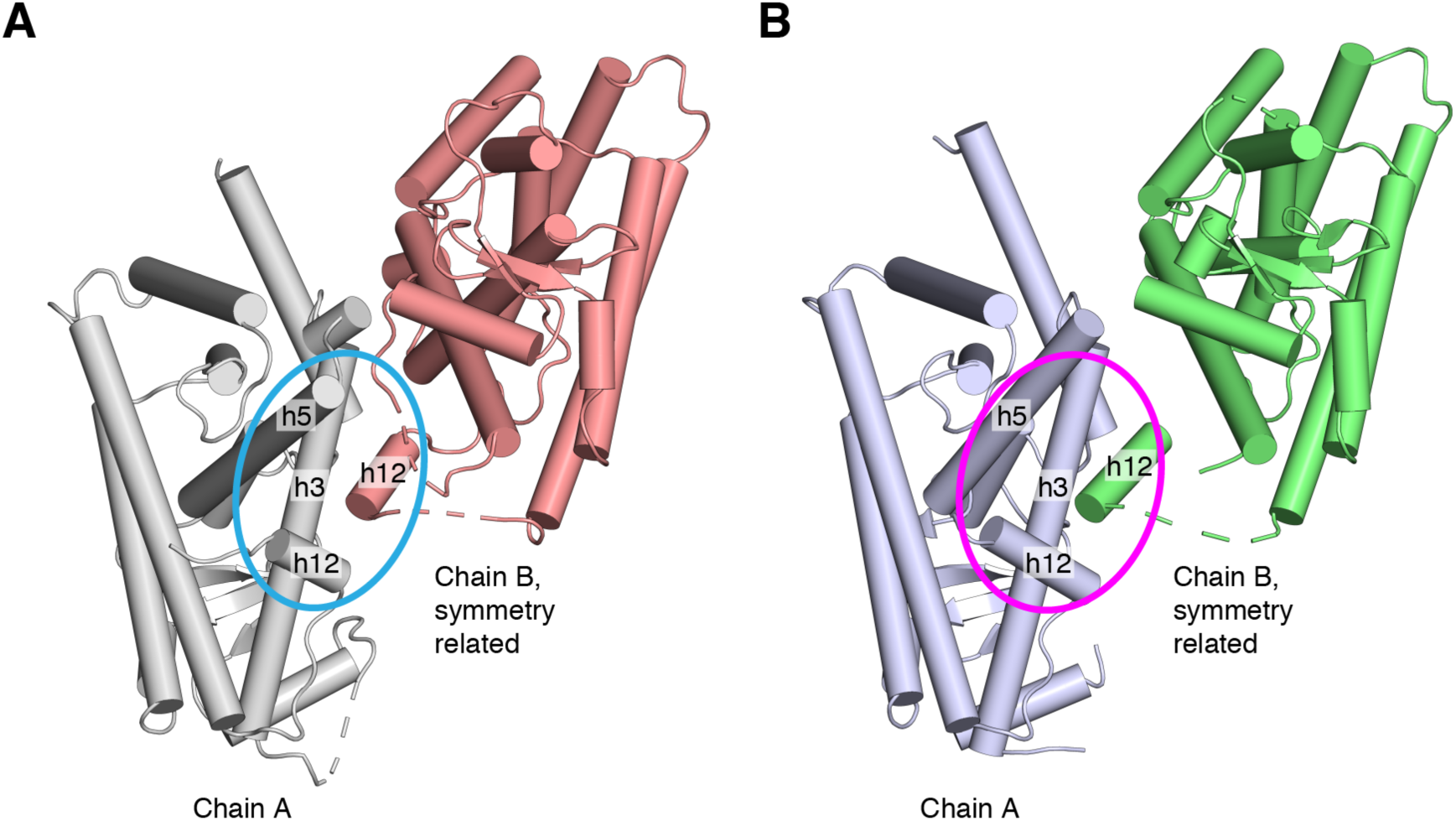
Distorted helix 12 conformations due to crystal contacts. Shown are chain A and the symmetry related chain B for the (**A**) T0070907-bound and (**B**) GW9662-bound (PDB 3B0R) PPARγ LBD crystal structures, where the chain B helix 12 docks into the AF-2 surface of chain A formed by helix 3, 5, and 12.

**Figure S3.**
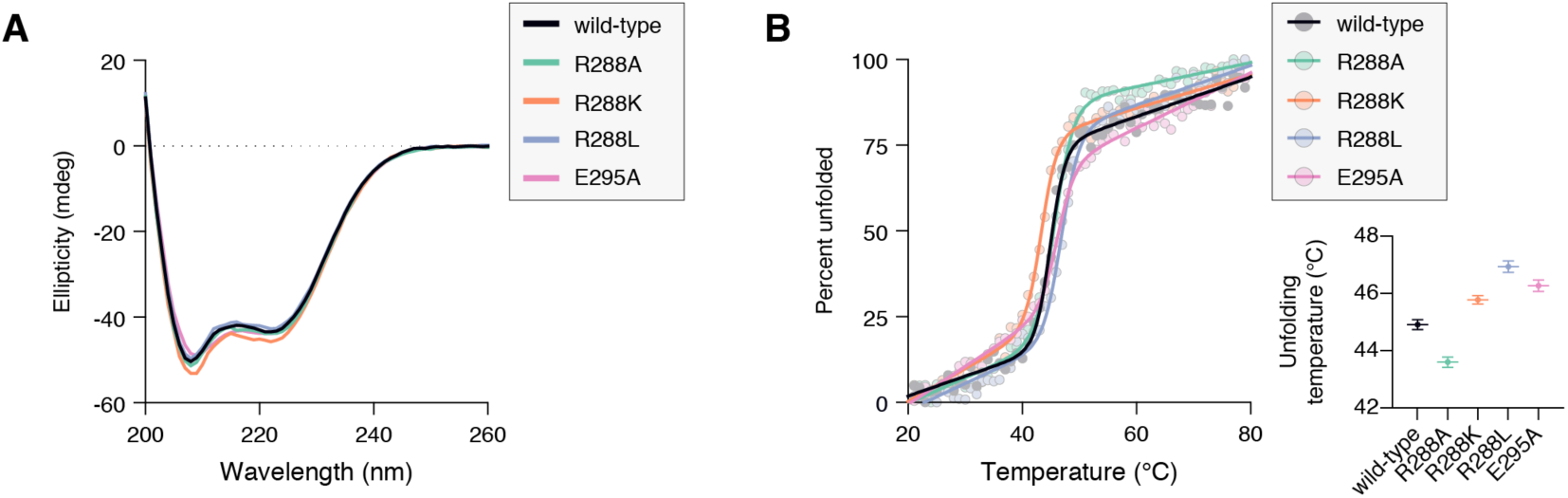
Circular dichroism (CD) spectroscopy data on PPARγ LBD mutants. (**A**) CD spectra and (**B**) CD thermal melt experiments (inset, fitted melting/unfolding temperatures).

**Figure S4.**
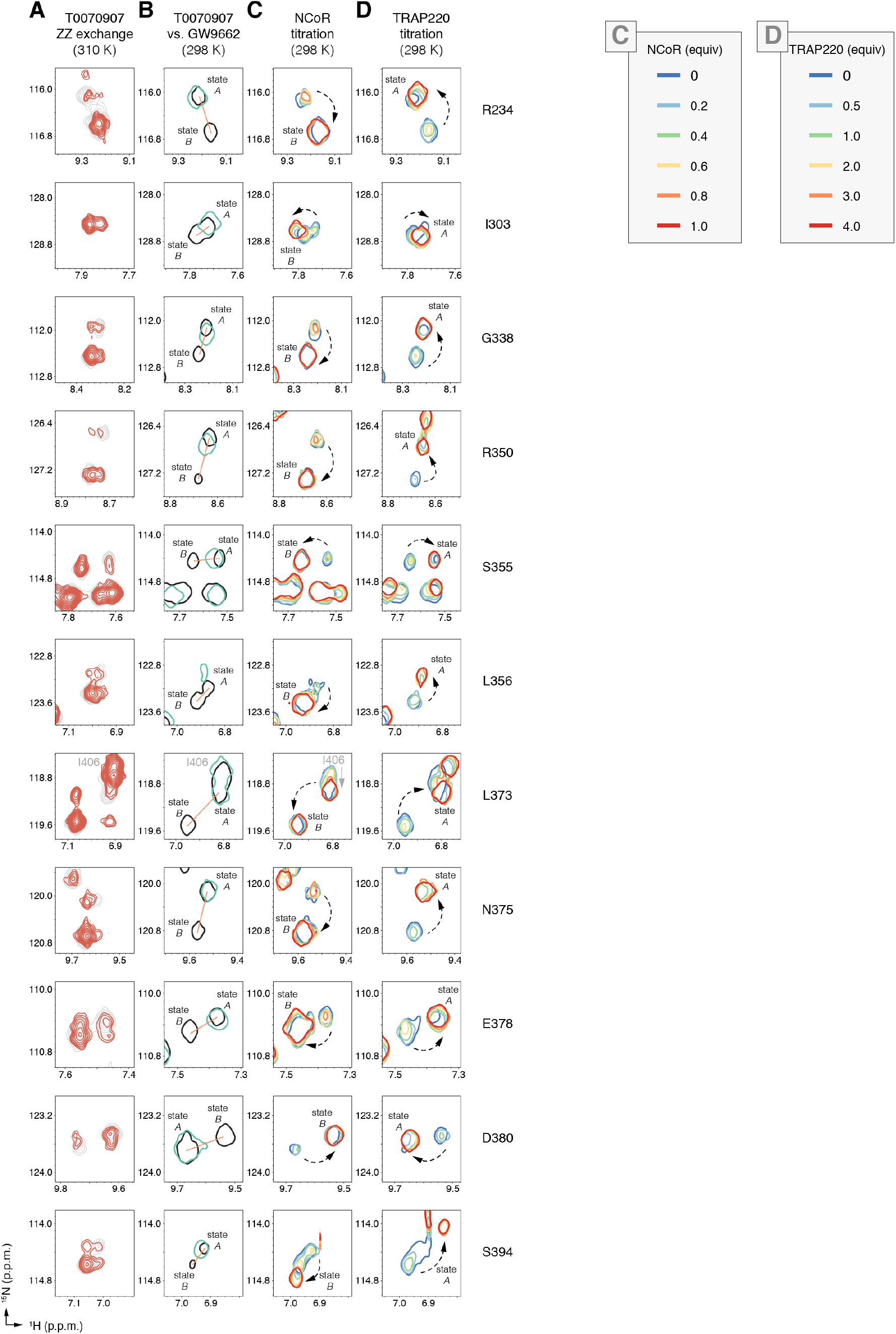
NMR reveals a slow global conformational change between two longlived conformations with distinct coregulator binding preferences. Data shown for the residues indicated to the right, which are displayed on the PPARγ LBD structure in Fig. 4b. (**A**) Snapshot overlays of ZZ-exchange ^15^N-HSQC NMR spectra of T0070907-bound ^15^N-PPARγ LBD; delay = 1 s (red peaks) and 0 s (grey peaks). (**B**) Snapshot overlays of [^1^H,^15^N]-TROSY-HSQC NMR spectra of ^15^N-PPARγ LBD bound to GW9662 (green) or T0070907 (black) shows that the single GW9662-bound G399 peak has similar chemical shift values to one of the two (connected by an orange line) T0070907-bound G399 peaks (state *A*); state *B* is uniquely populated by T0070907. (**C,D**) Snapshots of [^1^H,^15^N]-TROSYHSQC spectra of ^15^N-PPARγ LBD bound to T0070907 and titrated with (**C**) NCoR or (**D**) TRAP220 peptide.

